# Antibody repertoire and gene expression dynamics of diverse human B cell states during affinity maturation

**DOI:** 10.1101/2020.04.28.054775

**Authors:** Hamish W King, Nara Orban, John C Riches, Andrew J Clear, Gary Warnes, Sarah A Teichmann, Louisa K James

**Author notes:** To whom correspondence can be addressed, and.

## Abstract

In response to antigen challenge, B cells clonally expand, undergo selection and differentiate to produce mature B cell subsets and high affinity antibodies. However, the interplay between dynamic B cell states and their antibody-based selection is challenging to decipher in primary human tissue. We have applied an integrated analysis of bulk and single-cell antibody repertoires paired with single-cell transcriptomics of human B cells undergoing affinity maturation. We define unique gene expression and antibody repertoires of known and novel B cell states, including a pre-germinal centre state primed to undergo class switch recombination. We dissect antibody class-dependent gene expression of germinal centre and memory B cells to find that class switching prior to germinal centre entry dictates the capacity of B cells to undergo antibody-based selection and differentiate. Together, our analyses provide unprecedented resolution into the gene expression and selection dynamics that shape B cell-mediated immunity.

## Introduction

Effective immunity depends on the ability of B cells to evolve a functional antibody repertoire in response to antigen challenge. Following antigen encounter, activated B cells either differentiate into short-lived plasma cells or following cognate interaction with antigen-specific T cells can form germinal centres (GCs) within secondary lymphoid tissues, such as the spleen, peripheral lymph nodes and tonsils (Cyster and Allen, 2019). These GCs are transient structures in which B cells undergo iterative cycles of clonal expansion and somatic hypermutation in the variable regions of their immunoglobulin heavy and light chain genes followed by affinity-based selection for clones with high antigen-specificity. This highly dynamic process occurs in spatially and transcriptionally distinct dark and light zones (DZ and LZ) under the regulation of a network of specialised T follicular helper cells, follicular dendritic cells and macrophages (Mesin et al., 2016). B cells can differentiate and exit the GC reaction either as antibody-secreting plasmablasts committed to the plasma cell lineage or memory B cells, which are long-lived quiescent cells capable of being reactivated upon secondary exposure to the antigen (Suan et al., 2017b). The effector functions of antibodies expressed by B cells are broadly determined by antibody class (IgM, IgD, IgG, IgA, IgE) and more precisely by isotype or subclass (IgG1-4, IgA1-2), specified by the constant domain genes in the immunoglobulin heavy chain (IgH) locus. Although all naïve B cells express IgM and IgD, during maturation they may undergo class switch recombination, which involves the deletional recombination of IgM and IgD constant domain genes and expression of a different downstream constant domain gene (IgG1-4, IgA1-2 or IgE) (Stavnezer and Schrader, 2014). The combined outcomes of B cell differentiation, antigen affinity maturation and class switch recombination ultimately shape the antibody repertoire and the B cell-mediated immune response more broadly.

During their maturation in germinal centres, B cells express their antibody immunoglobulin genes as part of a membrane-bound complex termed the B cell receptor (BCR). Antigen-binding and downstream signalling of the BCR is a primary determinant of GC B cell survival and even differentiation through differential expression of key transcription factors regulated by BCR-mediated signalling (Kwak et al., 2019, Shlomchik et al., 2019). Studies have shown that BCR activation thresholds and downstream signalling can differ as a result of isotype-specific differences in the extracellular, transmembrane and intracellular domains of immunoglobulin proteins forming the BCR (Martin and Goodnow, 2002, Engels et al., 2014, Xu et al., 2014b). Thus, as well as shaping the effector functions of the subsequent antibody repertoire, class switch recombination contributes towards B cell survival or fate specification within the GC reaction. However, resolving the combined contribution of somatic hypermutation, maturation and class switching in the polyclonal context of primary human lymphoid tissues remains an enormous challenge for the field (Mesin et al., 2016). Furthermore, while it has long been held that class switch recombination occurs exclusively within the GC, as this is where the highest AICDA expression is detected, other work has demonstrated that class switch recombination often occurs prior to formation of the GC response (Roco et al., 2019, Toellner et al., 1996, Pape et al., 2003). This has raised questions about our current understanding of the cellular states and dynamics during human B cell maturation *in vivo* and demands a systematic and unbiased approach to better define how the human antibody repertoire is shaped through somatic hypermutation, class switch recombination and differentiation into different B cell fates.

We have applied an integrated strategy of bulk and single-cell antibody repertoire analysis paired with single-cell transcriptomics of human B cells from a model secondary lymphoid tissue, tonsils. We compare and contrast the antibody repertoires of major B cell subsets to reveal unique class switch hierarchies of memory B cells and plasmablasts. We then discover and define novel transcriptional B cell states during the GC response using single-cell RNA-seq. In particular, we reveal the unique gene expression of a B cell state primed to undergo class switch recombination before entering the GC. By leveraging the single-cell resolution of our datasets, we deconvolve the contribution of somatic hypermutation and antibody class to gene expression patterns linked with altered BCR signalling, B cell maturation and fate decisions within the GC. Finally, we define diverse memory B cell states within secondary lymphoid tissue and explore the impact of class switch recombination on their functional potential. Our analyses reveal a striking importance for class switch recombination in shaping B cell fate and maturation in the GC and memory B cell fate that reframes our understanding of antibody-based selection and B cell differentiation.

## Results

### Subset- and subclass-specific antibody class switch recombination landscapes

To begin to untangle the antibody-based selection of different B cell fates, we characterised the antibody repertoires of four broadly defined B cell subsets from the human tonsil; naïve, germinal centre (GC) B cells, memory B cells (MBCs) and plasmablasts, in addition to total CD19+ B cells (Figure 1A). We applied a subclass-specific and quantitative unique molecular identifier-based repertoire sequencing protocol for IgH VDJ sequences (Horns et al., 2016). Naïve B cells were comprised of more than 95% unswitched and unmutated IgM and IgD sequences, while GC and MBC samples consisted of both switched and unswitched IgH sequences with elevated somatic hypermutation (Figure 1B-C). Plasmablasts were nearly all switched and highly mutated. Consistent with the low abundance of IgE^+^ B cells in human tonsils, we detected only a single IgE sequence, preventing us from drawing any conclusions about IgE-expressing B cells.

**Figure 1.**
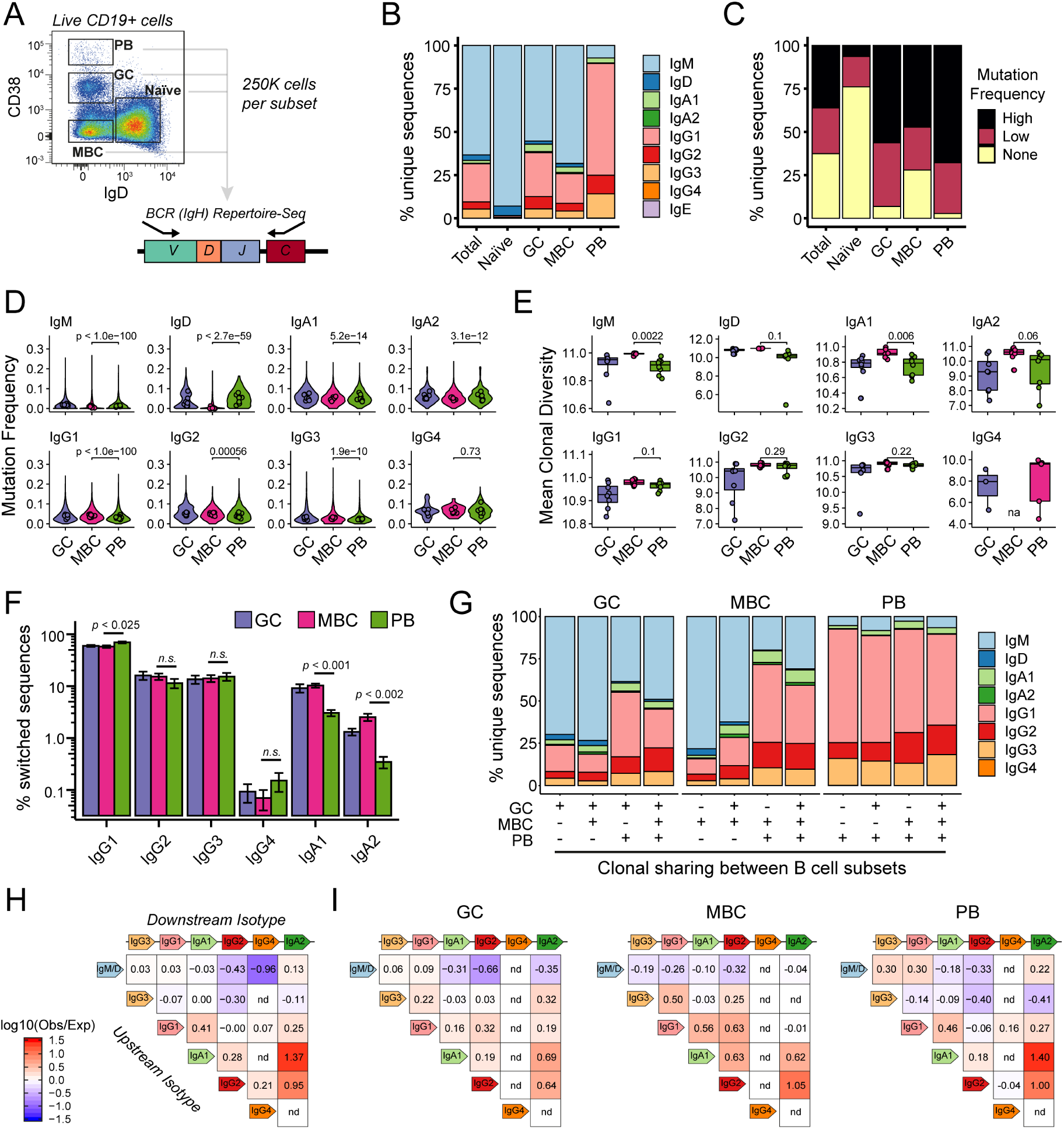
Subclass- and subset-specific features of human tonsillar B cell repertoires. A) Cell sorting strategy to isolate naïve, germinal centre (GC), memory B cells (MBC) and plasmablasts (PB) from live CD19+ human tonsillar B cells for antibody repertoire analysis. Representative of *n*=8. B) Mean antibody subclass frequencies within each B cell subset across donors. C) Mean frequencies of antibody somatic hypermutation levels (None, Low, High) within each B cell subset across donors. D) Somatic hypermutation frequencies for subclass-specific antibody sequences within each B cell subset. Violin plots show all unique sequences per subset, while points represent mean for each donor. E) Mean clonal diversity scores per donor of subclass- and subset-specific B cell clones. F) Mean switched antibody subclass frequencies within each B cell subset across donors. G) Mean subclass frequencies for expanded clones spanning different B cell populations (x axis). For each class of clone, subset-specific members are examined (see groups at top). H) Observed/expected frequencies for isotype pairs detected in reconstructed phylogenies of clonally-related sequences. Antibody subclasses are ordered according to the IgH locus. nd denotes not detected. I) Same as in H), except phylogenies are restricted to subset-specific sequences.

Higher IgH somatic hypermutation frequencies within the GC are typically a reflection of higher affinity BCRs and are proposed to bias GC B cells towards the plasmablast cell fate rather than the MBC fate (Shinnakasu et al., 2016, Suan et al., 2017a). In keeping with this, plasmablast-derived antibody repertoires generally contained higher somatic hypermutation frequencies than those of MBCs at a bulk level (Figure 1C). However, by resolving for antibody subclass, we found that while MBC-derived IgM and IgD sequences were consistently less mutated than plasmablast-derived IgM and IgD, the somatic hypermutation levels for switched isotypes were broadly similar between different B cell subsets (Figure 1D). Comparison of the clonal diversity of subclass-specific MBCs and plasmablasts revealed unswitched and IgA^+^ MBCs were less clonally expanded (as evidenced by higher diversity) than plasmablasts of the same isotype, while IgG^+^ MBCs and plasmablasts appeared to have clonally expanded to similar degrees (Figure 1E). This is not likely explained by differences in their somatic hypermutation frequencies (Figure 1D), but instead may reflect differential selection or differentiation processes linked with specific class switch recombination outcomes.

We next examined whether there were differences in antibody subclass frequencies or the manner in which subclasses might have arisen in our broadly defined B cell populations. Intriguingly, we observed that as well as an increased propensity to retain IgM expression (Figure 1B), MBCs were 3.3- or 7.3-fold more likely than plasmablasts to express IgA1 or IgA2 respectively, while plasmablasts were significantly more likely to express IgG1 (Figure 1F). These enrichments were linked with specific B cell fates even for expanded clones spanning different B cell populations (Figure 1G). Finally, to explore how these unique class switch patterns might have arisen, we reconstructed phylogenies for 28,845 expanded B cell clones and calculated the likelihood that specific class switch recombination events were observed compared to that expected by chance (Figure 1H-I). Clonal lineages within the MBC pool exhibited greater likelihoods for switching of isotype pairs located close to each other in linear space along the IgH locus, compared to plasmablast clones which demonstrated a more eclectic pattern of class-switch likelihoods (Figure 1I). Of note, both the antibody subclass frequencies and reconstructed class switch hierarchies of MBCs closely resembled those of GC cells, consistent with models that propose a stochastic exit of MBCs from the GC (Duffy et al., 2012, Good-Jacobson and Shlomchik, 2010). Together, these analyses reveal that the antibody-based selection mechanisms of two major mature B cell subsets (MBCs and plasmablasts) exhibit important differences related primarily to their propensity to have undergone class switch recombination earlier in their maturation.

### Single-cell atlas of tonsillar immune cells defines B cell states during affinity maturation

To better understand the antibody-based selection mechanisms shaping the maturation of different B cell subsets, we performed single-cell RNA-seq (scRNA-seq) paired with single-cell B cell repertoire VDJ sequencing (scVDJ-seq) for unsorted tonsillar immune cells from the same samples used for our bulk B cell repertoire analyses (Figure 2A-C). After stringent quality control, we retained the transcriptomes of 32,607 immune cells (*n*=7; median of 3142 and mean of 4658 cells per donor) from which we identified 30 transcriptionally distinct cell types (Figure 2D-F). Although our primary focus was understanding B cell maturation, given the importance of other immune cell populations in this process we also annotated varied T cell and non-lymphoid populations, including naïve and/or central memory (NCM) CD4^+^ and CD8^+^ T cells, follicular helper (TfH) T cells, follicular regulatory T (Tfr) cells, regulatory T cells (Treg), cytotoxic CD8^+^ T cells, innate lymphoid cells (ILCs), natural killer (NK) cells, follicular dendritic cell (FDC), plasmacytoid-derived dendritic cell (pDCs), classical dendritic cell (cDC1), and several macrophage (MAC) clusters (Figures 2D-E and S1) that will act as a valuable resource for those studying immune cell dynamics in human secondary lymphoid tissues.

**Figure 2.**
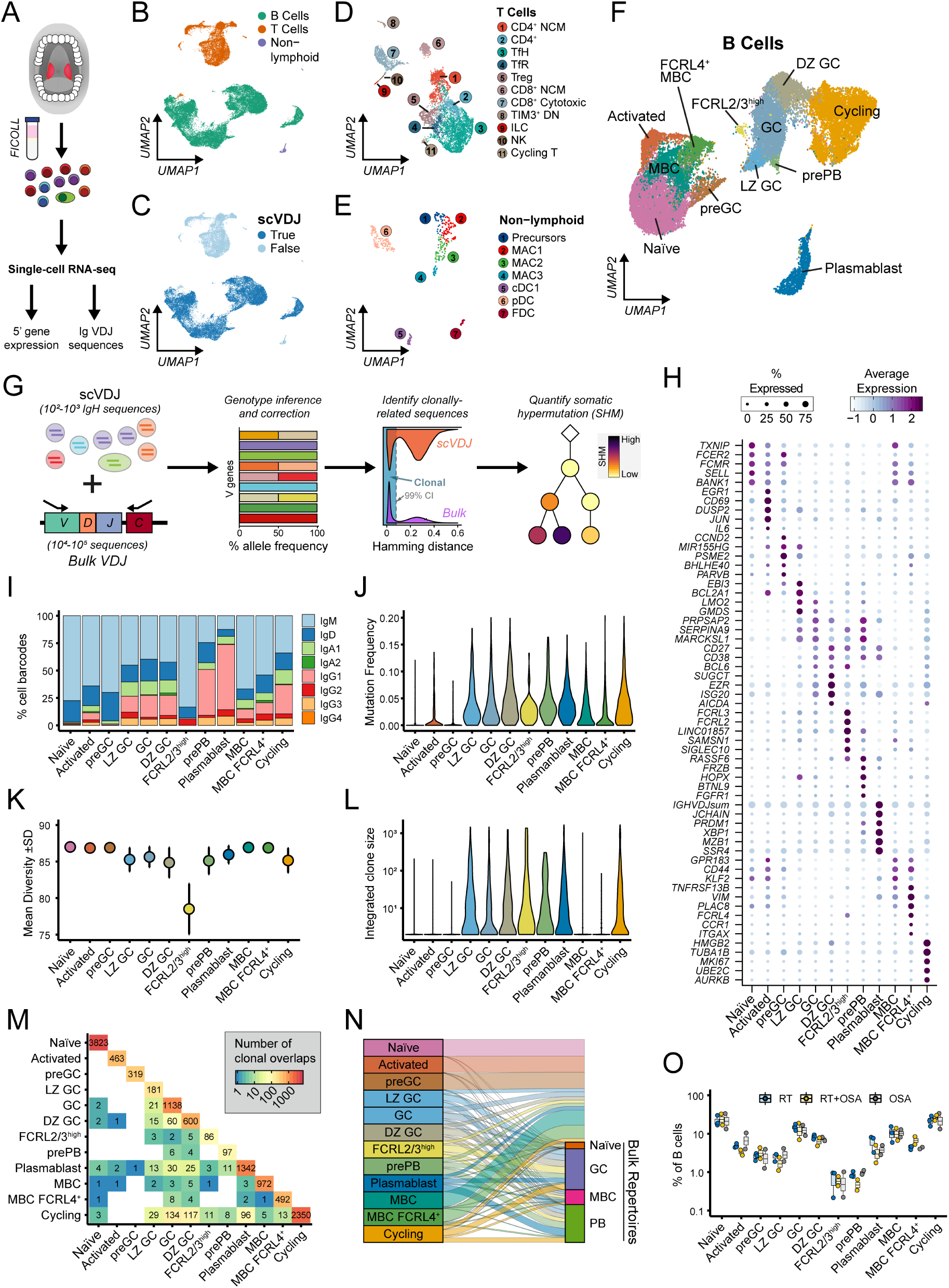
A single-cell atlas of human tonsillar immune cells to understand B cell maturation. A) Schematic of tonsillar immune cell isolation followed by single-cell profiling of gene expression and antibody sequences. B) UMAP projection of tonsillar immune scRNA-seq data (32,607 cells; 7 donors) annotated as B, T or non-lymphoid cells. C) Same as in B), with cells annotated for whether a high quality immunoglobulin scVDJ sequence was assembled. D) UMAP projection of T cell populations in the tonsillar immune scRNA-seq data, including CD4^+^ naïve or central memory (CD4^+^ NCM), CD4^+^, T follicular helper (TfH), T follicular regulatory (TfH), T regulatory (Treg), CD8^+^ naïve or central memory (CD8^+^ NCM), CD8^+^ cytotoxic, TIM3^+^ CD4/CD8 double-negative (TIM3^+^ DN) and cycling T cells, in addition to innate lymphoid cells (ILC) and natural killer (NK) cells. E) UMAP projection of non-lymphoid cell populations in the tonsillar immune scRNA-seq data, including monocyte/macrophage precursors (Precursors), macrophages (MAC1, MAC2, MAC3), conventional dendritic cell 1 (cDC1), plasmacytoid-derived dendritic cell (pDC) and follicular dendritic cell (FDC) subsets. F) UMAP projection of B cell populations in the tonsillar immune scRNA-seq data, including naïve, activated, pre-germinal centre (preGC), light zone GC (LZ GC), GC, dark zone (DZ GC), cycling, FCRL2/3^high^ GC, pre-plasmablasts (prePB), plasmablasts, MBC and FCRL4^+^ MBC subsets. G) Schematic of scVDJ and bulk repertoire integration, including genotype inference and correction, identification of clonally-related sequences and quantitation of somatic hypermutation. H) Mean expression of key marker genes used to define B cell scRNA-seq clusters. Frequency of cells for which each gene is detected is denoted by size of the dots. I) Relative scVDJ-derived antibody subclass frequencies in different B cell states. J) Somatic hypermutation frequencies of scVDJ-derived antibody genes in different B cell states. K) Clonal diversity scores of B cell clones identified in scRNA-seq dataset. Error bars denote ±SD. L) Number of members per clonotype in different B cell states from integrated repertoire analysis. M) Co-occurrence of expanded clones across B cell states. Numbers reflect a binary detection event. N) Clonal relationships between scRNA-seq-defined B cell states and repertoires of sorted B cell subsets. O) Relative frequencies of different B cell states separated by clinical indication for tonsillectomy. OSA; obstructive sleep apnoea (*n*=2), RT; recurrent tonsillitis (*n*=3), RT+OSA (*n*=2).

We next characterised tonsillar B cell states using both their unique gene expression and scVDJ-derived antibody repertoire features. To improve the power and accuracy of our scVDJ analyses we applied a novel strategy of integrating single-cell and bulk repertoires to benefit from the deeper sampling of IgH sequences from bulk repertoires to enhance genotype correction, identification of clonally-related sequences and quantitation of somatic hypermutation levels for scVDJ-derived sequences (Figure 2G). We defined gene expression signatures for 12 distinct B cell types or states (Figure 2H) and complemented marker gene-based annotation with antibody isotype frequencies (Figure 2I), somatic hypermutation levels (Figure 2J), clonal diversity (Figure 2K-L) and relationships with other B cell subsets (Figure 2M-N). We identified all major stages of B cell maturation, including naïve, activated, GC (including both LZ and non-proliferating DZ cells), MBCs, tissue-resident FCRL4+ MBCs, plasmablasts and a cycling population consisting mostly of DZ GC cells (Figure 2H). Whilst apoptotic cells normally comprise a sizeable proportion of GC B cells these were not retained in our analysis as they failed to generate sufficiently high quality transcriptomic data. Our analysis of B cell states also identified a “preGC” B cell state expressing unmutated IgM and IgD that transcriptionally shared markers with both naïve and LZ GC populations, but had yet to acquire features consistent with B cell maturation in the GC such as *CD27* and *CD38* expression, hypermutated antibody genes or clonal expansion (see Figure 3 for further details). We also annotated a population of class switched and hypermutated GC B cells that amongst other transcriptionally unique features (*FRZB, BTNL9, FGFR1*) express low to intermediate levels of the plasmablast-specific transcription factors *PRDM1* and *XBP1* (Figure 2H), suggesting that these cells may be a pre-plasmablast (prePB) state within the GC. Finally, we discover a transcriptionally distinct and clonally-expanded IgM^+^ B cell population in the GC with elevated expression of genes associated with inhibitory BCR signalling, such as *FCRL2, FCRL3, SAMSN1*, and *SIGLEC10* that we have labelled as FCRL2/3^high^ GC B cells. Some of these cells were part of large expanded GC-derived clones that also contained MBCs or plasmablasts (Figure 2M-N), indicating that this cell state arises as part of productive GC reactions and is unlikely to be derived from a separate B cell lineage.

**Figure 3.**
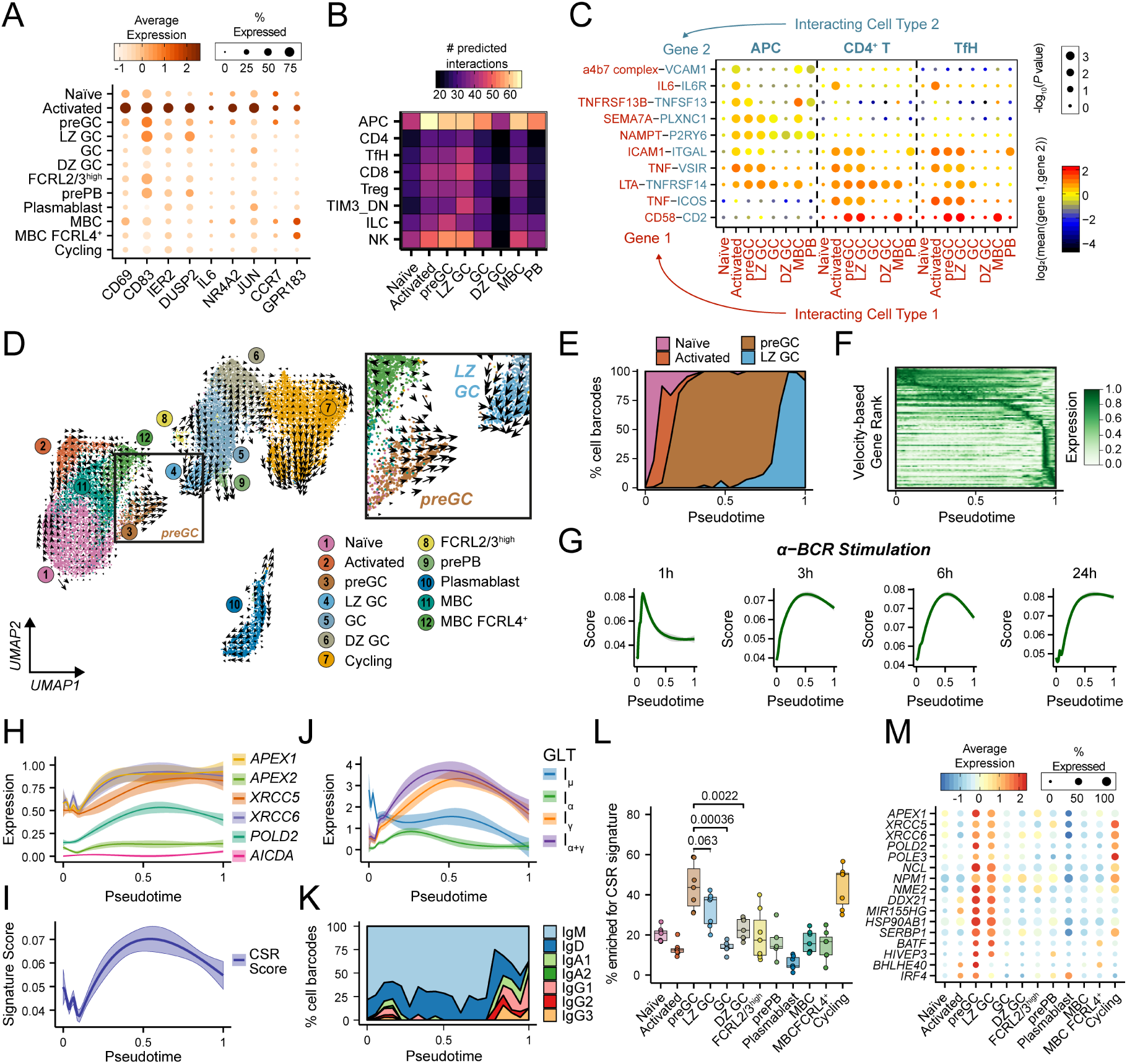
Reconstruction of *in vivo* B cell activation reveals class switch recombination in a preGC state. A) Mean expression of key marker genes for activated B cells in scRNA-seq. B) Frequency of significant predicted ligand-receptor pair interactions between major B cell states and different T cell populations and antigen-presenting cells (APCs) performed using CellPhoneDB. C) Selected ligand–receptor interactions between B cell subsets and APCs, CD4+ T cells and TfH cells with CellPhoneDB. D) Grid-based visualisation of tonsillar B cell RNA velocities. Arrow size conveys strength of predicted directionality. E) Relative frequencies of B cell types in a velocity-based pseudotime reconstruction of B cell activation and GC formation. F) Heatmap depicting dynamic gene expression across velocity-based pseudotime reconstruction in E). G) Smoothed anti (α)-IgM-treatment gene signature scores (±95% CI) across velocity-based pseudotime. H) Smoothed expression of class switch recombination genes differentially expressed through velocity-based pseudotime. I) Smoothed class switch recombination gene signature score through velocity-based pseudotime. J) Smoothed expression of I promoter germline transcripts for IgM (I_µ_), IgA (I_α_) and IgG (I_γ_) through velocity-based pseudotime. I_α+γ_ denotes the sum of I_α_ and I_γ_ expression. K) Relative antibody subclass frequencies across velocity-based pseudotime. L) Relative frequencies of cells with high class switch recombination signature scores in different B cell states for each donor (*n* = 7). *p* values denote results of Student’s T test. M) Mean expression of genes implicated in class switch recombination for B cell subsets.

Crucially, all annotated B cell states were observed at reproducible frequencies across patients, regardless of their history of tonsillitis (Figure 2O), although exactly how some of these transcriptional states relate to other human B cell populations will require further work. We also note that we have identified a greater diversity of B cell fates in our paediatric tonsil samples (typically <10 years old) than previous single-cell studies of other human lymphoid tissues from adult donors (typically >40 years old) (Madissoon et al., 2019, James et al., 2020), highlighting the importance of profiling immunologically active tissues to understand B cell maturation. Together, our uniquely comprehensive transcriptomic and repertoire analyses of a model secondary lymphoid organ have allowed us to create a detailed overview of human B cell maturation that will allow us to interrogate gene expression and antibody repertoire dynamics before, during and after maturation in the GC.

### Dynamic gene expression during human B cell activation and GC formation

Initially in a B cell response, B cells acquire antigen, either in soluble form or displayed on the surface of follicular antigen-presenting cells (APCs), which results in their activation and migration from the follicle to the T cell zone where they can form or participate in GC reactions. However, reconstructing the events during this important process in primary human tissues is extremely challenging. We therefore used our single-cell profiling of human B cell maturation to explore the dynamics of early B cell activation and GC formation. We first identified an activated B cell state with elevated expression of known activation marker genes (Activated; *CD69, CD83, JUN*) (Figures 2 and 3A) and which demonstrate the highest frequency of predicted cell-cell communication with APCs (FDCs, other dendritic cells, macrophages) (Figure 3B). This activated B cell state appeared capable of coordinating help from both APCs and T cells through IL6 signalling (Arkatkar et al., 2017, Ise et al., 2018) and/or ICAM1-ITGAL1 (LFA-1) interactions (Zaretsky et al., 2017) (Figure 3C). Many of these same predicted cell-cell interactions were also detected in preGC B cells (Figure 3C), suggesting that these might reflect a transitional cell state between antigen dependent-activation and GC entry or formation. Indeed, we found that preGC B cells exhibit a strong directionality towards the light zone of the GC using RNA velocity inference (Figure 3D). We next reconstructed a pseudotemporal trajectory of naïve, activated, preGC and LZ GC B cells, revealing a continuum of gene expression from early activation events to *bona fide* GC B cells and allowing us to identify dynamic expression of key signalling molecules and transcription factors (Figures 3E-F and S2A). Crucially, these gene expression changes through pseudotime were well correlated with an experimentally-derived time course of *in vitro*-stimulated B cells (Figure 3G; Shinohara et al., 2014), strongly supporting our pseudotemporal ordering of defined B cell states during B cell activation and GC entry. This roadmap of B cell activation may allow future improvement to *in vitro* B cell culture protocols to better model B cell activation dynamics *in vivo*.

One surprising result from our reconstruction of human B cell activation and GC entry was the discovery that several genes associated with class switch recombination were most highly expressed prior to GC entry, with a specific enrichment of these genes in the preGC B cell population (Figure 3H-I). This included *APEX1* expression, of which its translated product APE1 is required for class switch recombination to occur in a dose-dependent manner (Masani et al., 2013, Xu et al., 2014a) and is expressed almost exclusively by non-GC B cells (Figure S2B; Roco et al., 2019). We also found that preGC B cells had the highest expression of IgH germline transcripts (GLTs) (Figure 3J), which preceded switching from IgM/IgD to other isotypes (Figure 3K), in fitting with a recent study describing GLT transcription prior to GC formation in mouse models (Roco et al., 2019). These findings contrast with the prevailing view that class switch recombination occurs in the DZ of the GC, where AICDA expression is highest. However, we detected little enrichment of our class switch recombination signature in non-cycling DZ GC B cells (Figure 3L), and although there was an enrichment of class switch recombination genes in cycling B cell populations, this likely reflects their involvement in cell cycle-linked DNA recombination and repair. Our analysis also highlights many other genes previously linked with class switch recombination (Figure 3M), including those capable of binding to switch region sequences within the IgH locus (*NME2, NCL, DDX21*), interacting with the class switch recombination machinery (*NPM1, SERBP1*) or regulating *AICDA*/AICDA transcript/protein stability (*mir155HG, HSP90AB1*) (Borggrefe et al., 1998, Hanakahi et al., 1997, Mondal et al., 2016, Orthwein et al., 2010, Shinozaki et al., 2006, McRae et al., 2017, Zheng et al., 2019). Notably, the microRNA gene *miR155HG*, formerly called BIC (B-cell Integration Cluster), was up-regulated in preGC B cells and has been shown to be essential for B cells to form GCs and undergo class switch recombination in mice (Thai et al., 2007, Vigorito et al., 2007). Furthermore, the transcription factor genes *BATF, IRF4* and *BHLHE40* are enriched in the preGC B cell state, of which BATF and IRF4 are known to regulate GC formation in a B cell-intrinsic manner (Morman et al., 2018, Willis et al., 2014). Intriguingly, BHLHE40 is capable of binding to the major regulatory regions α1 RR and α_2_ RR of the IgH locus (Figure S2C-D), implicating this poorly understood transcription factor in the regulation of class switch recombination prior to GC formation. Together, these analyses strongly support a model where class switch recombination occurs primarily before formation of the GC response (Roco et al., 2019, Toellner et al., 1996, Pape et al., 2003) and our detailed gene expression analyses define the cellular state involved.

### Antibody-based selection of B cell fate in the germinal centre

Class switch recombination before entry into the GC has the potential to dramatically influence the antibody-based selection of B cells within the subsequent GC reaction as a consequence of differential signalling through the membrane-bound immunoglobulin BCR. We therefore turned to dissect the gene expression dynamics linked with antibody-based selection and fate specification of B cells within the GC reaction.

During affinity maturation and selection in the GC, B cells cycle between physically distinct light and dark zones. While we clearly identified LZ and DZ B cell populations in our scRNA-seq dataset, we also found that many GC B cells existed in a continuum between these two states (Figure 4A-B) similar to previous reports (Milpied et al., 2018), with the exception of the FCRL2/3^high^ and prePB clusters which existed as transcriptionally distinct states (Figures 2H and 4C). In addition to unique gene expression patterns, these two sub-populations of GC B cells also exhibit unique patterns of class switching, with prePB B cells in the GC more likely to express class-switched isotypes and FCRL2/3^high^ GC B cells more likely to retain expression of IgM and IgD (Figures 2I and 4D). Furthermore, cells clonally related to FCRL2/3^high^ GC B cells were almost exclusively IgM^+^ and had rarely undergone class-switching (Figure 4E). These observations suggested that the outcome of class switch recombination may be linked with specific gene expression programs of GC B cells.

**Figure 4.**
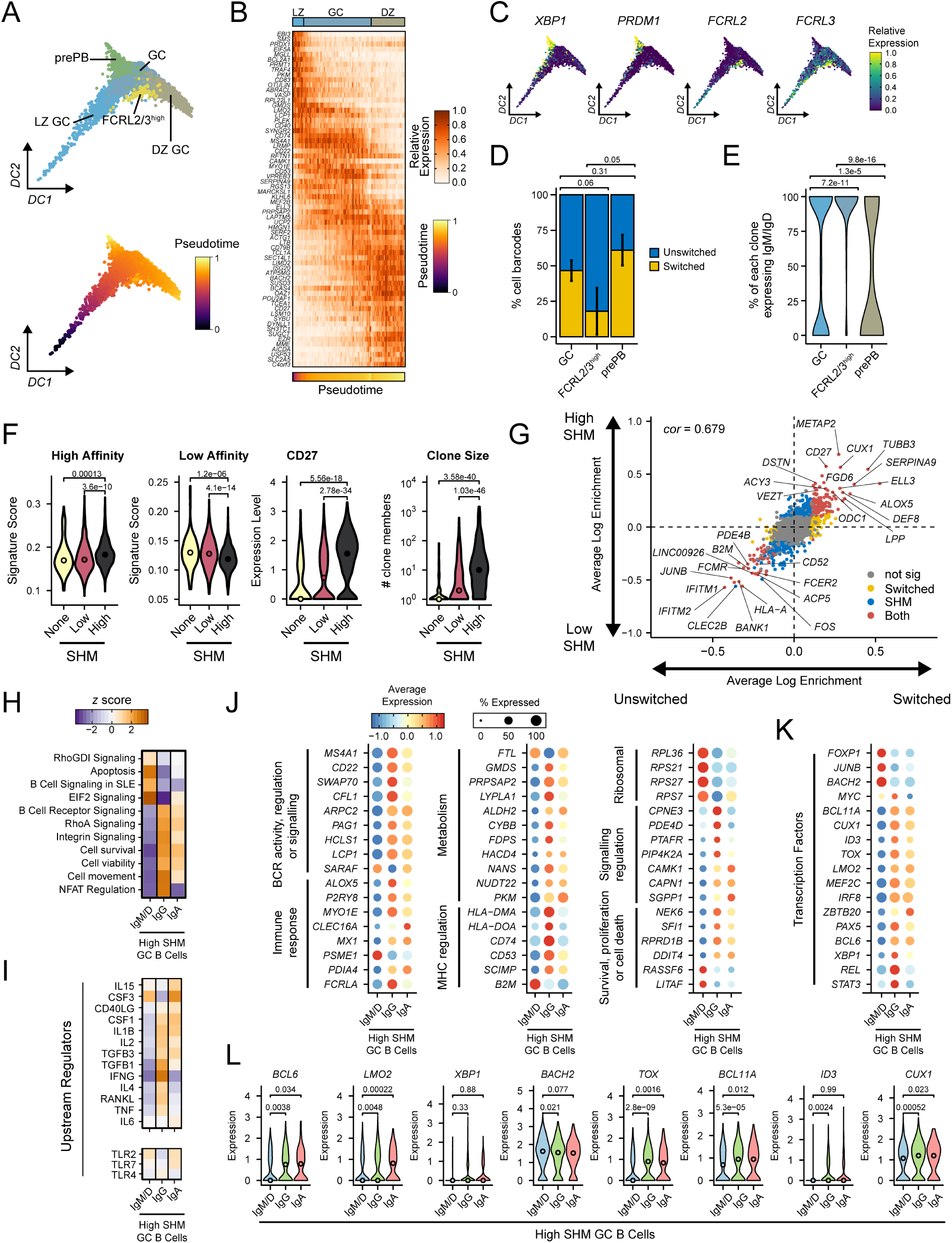
Influence of antibody class on B cell fate and function within the germinal centre. A) Diffusion-based graph visualisation and pseudotemporal ordering of GC B cell scRNA-seq populations. B) Single-cell gene expression heatmap of major GC B cell states ordered by pseudotime within each cluster. C) Expression of key marker genes for prePB and FCRL2/3^high^ GC B cells. D) Relative frequency of GC B cells that have undergone class switch recombination (*n* = 6). “GC” is the combination of LZ GC, GC and DZ GC clusters in (A). Error bars denote SEM. E) Percentage of members within each GC B cell clone that express IgM or IgD for clones that contain FCRL2/3^high^ or prePB cells. Clonal families include sequences from both scVDJ and bulk repertoires. F) High and low affinity gene signature scores for GC B cells (LZ GC, GC and DZ GC clusters) grouped by antibody somatic hypermutation (SHM) frequency (*n =* 2045 cells). Also shown are *CD27* expression levels and integrated clone sizes. G) Scatterplot comparing log enrichment of genes in class-switched vs unswitched (x axis) and high vs low affinity/SHM (y axis) GC B cells. Colour denotes statistical significance and c*or* denotes Pearson’s Correlation coefficient. H) Heatmap of significantly enriched gene ontologies for genes enriched in class-specific high affinity/SHM GC B cells. I) Same as in (H), but for gene pathways downstream of cytokines (upper panel) and toll-like receptors (TLRs). J) Mean expression of genes enriched in class-specific high affinity/SHM GC B cells grouped by predicted functions. K) Mean expression of transcription factor genes enriched in class-specific high affinity/SHM GC B cells. L) Single-cell gene expression of transcription factors in class-specific high affinity/SHM GC B cells.

To address this, we first needed to examine the contribution of antibody maturation state to GC B cell gene expression programs, given lower average somatic hypermutation frequencies of IgM^+^ B cells compared to B cells expressing switched isotypes (Figure 1D). We leveraged our paired single-cell VDJ and transcriptomic datasets to stratify all non-cycling GC B cells (excluding prePB and FCRL2/3^high^ populations) based on their IgH somatic hypermutation frequencies as a proxy for affinity (Figure 4F). Strikingly, GC B cells with high or low somatic hypermutation were significantly enriched with gene sets derived from experimentally determined high- or low-affinity antigen-binding B cells respectively (Shinnakasu et al., 2016), higher expression of the B cell maturation marker CD27 and larger clone sizes (Figure 4F), reflecting increased expansion and maturation based on BCR affinity. We found that the direct comparison of GC B cells expressing different antibody classes was confounded by gene expression linked with high and low somatic hypermutation frequencies (Figure 4G), consistent with high and low affinity binding events differentially regulating GC B cells (Shinnakasu et al., 2016). To overcome this, we examined GC B cells with matched somatic hypermutation levels expressing different antibody classes (IgM/D, IgG or IgA) at single-cell resolution. This revealed differential expression of genes involved directly in cell survival, BCR signalling, antigen presentation, immune responses and metabolism, as well as other pathways more indirectly linked with BCR activity (RhoA/RhoGDI, Integrin, NFAT, eIF2), between unswitched and switched GC B cells (Tybulewicz and Henderson, 2009, Mielke et al., 2011, Arana et al., 2008, Scharenberg et al., 2007) (Figure 4H-J). These differences may be linked with differential exposure to T cell-derived cytokines such as IL4, TGFB, IFNG and CD40LG, or signalling through different toll-like receptors (TLR) (Figure 4I). Intriguingly, several genes involved with GC confinement or regulating B cell niche homing were up-regulated in IgG^+^ and IgA^+^ GC B cells, such as genes required for CXCL12-mediated migration to GCs (*LCP1* and *MYO1E* (Dubovsky et al., 2013, Girón-Pérez et al., 2020)) and the GC confinement receptor *P2RY8* (Muppidi et al., 2014).

Although most gene expression differences were comparable between IgG and IgA and we identified few significant or meaningful differences for subclass-specific B cells (Figure S3), one interesting example of class-specific gene expression was the enrichment of *CLEC16A* in IgA^+^ GC B cells given that this gene is associated with a selective IgA immunodeficiency (Ferreira et al., 2010). Finally, to try and understand the upstream regulation of these class-specific gene expression networks we examined the expression of transcription factors within class-specific GC B cells (Figure 4K-L). We found that IgM^+^ B cells express lower levels of transcription factors like *BCL6, XBP1* and *ID3* known to regulate the ability of B cells to remain in the GC or differentiate but higher levels of the transcription factor *BACH2* that represses plasma cell differentiation (Todd et al., 2009, Gloury et al., 2016, Huang et al., 2014, Shinnakasu et al., 2016). We also found differential expression of other transcription factors such as *LMO2, TOX, BCL11A* and *CUX1*, suggesting that they may play a role in the unique transcriptional wiring of switched and unswitched B cells within the GC. Our single-cell resolution of the GC B cell response has allowed us to uncouple antibody affinity and class and to dissect the differential contributions of these two critical arms of the B cell repertoire in shaping B cell fate and function in the GC. Importantly, these differential gene expression patterns suggest varying abilities of switched and unswitched B cells to survive and reside in the GC and establish that one of the dominant influences in shaping antibody-based selection in the GC is whether a B cell has undergone class switch recombination.

### Diverse memory B cell states and activation dynamics in secondary lymphoid tissue

Maturation state and antibody class appeared to both impact gene expression dynamics within the GC, presumably through membrane-specific isoforms of immunoglobulin as part of the BCR. Upon GC exit and differentiation, plasmablasts lose membrane-bound immunoglobulin expression and instead start to secrete large amounts of soluble antibody. In contrast MBCs retain BCR expression of which the antibody isotype may influence the phenotypic properties of different MBC subsets (Engels and Wienands, 2018).

We therefore sought to determine whether antibody class expression by MBCs might be linked with different functional abilities and to better define the heterogeneity within the MBC pool in secondary lymphoid tissue. A significant proportion of memory B cells are unswitched (Figure 1B), so to examine potential differential gene expression across class switched memory B cells subsets we generated paired single-cell transcriptomics and VDJ repertoires for IgD-depleted or IgM/IgD-depleted MBCs from the same tonsillar immune cell preparations analysed previously. Dataset integration and quality control provided 21,595 high-quality MBC single-cell transcriptomes that we annotated with 11 clusters reflecting different MBC subsets and states (Figure 5A-C), all of which lacked marker gene expression for naïve or GC B cells (Figure S4A). In addition to tissue-resident FCRL4^+^ MBCs previously identified (Figure 2; Ehrhardt et al., 2005), we annotated two rare *CR2*/CD21^low^ MBC subsets resembling populations described elsewhere (Lau et al., 2017, Thorarinsdottir et al., 2016) (Figures 5A,C and S4B). However, the majority of MBC diversity within human tonsils appeared to reflect differences in cellular state or signalling activity rather than distinct cell types, such as different activation states (Activated 1 and Activated 2), heat shock protein (HSP)-related gene activity (HSP-response), and an IFN-responsive MBC state (IFN-response) (Figures 5C and S4C). We also identified a preGC MBC population with similar gene expression to naïve preGC cells and an enrichment for class switch recombination genes (Figures 5C and S4D-E), as well as an FCRL2/3^high^ MBC state similar to the FCRL2/3^high^ GC population (Figures 5C and S4D), suggesting that these may be widely shared functional states spanning multiple B cell fates.

**Figure 5.**
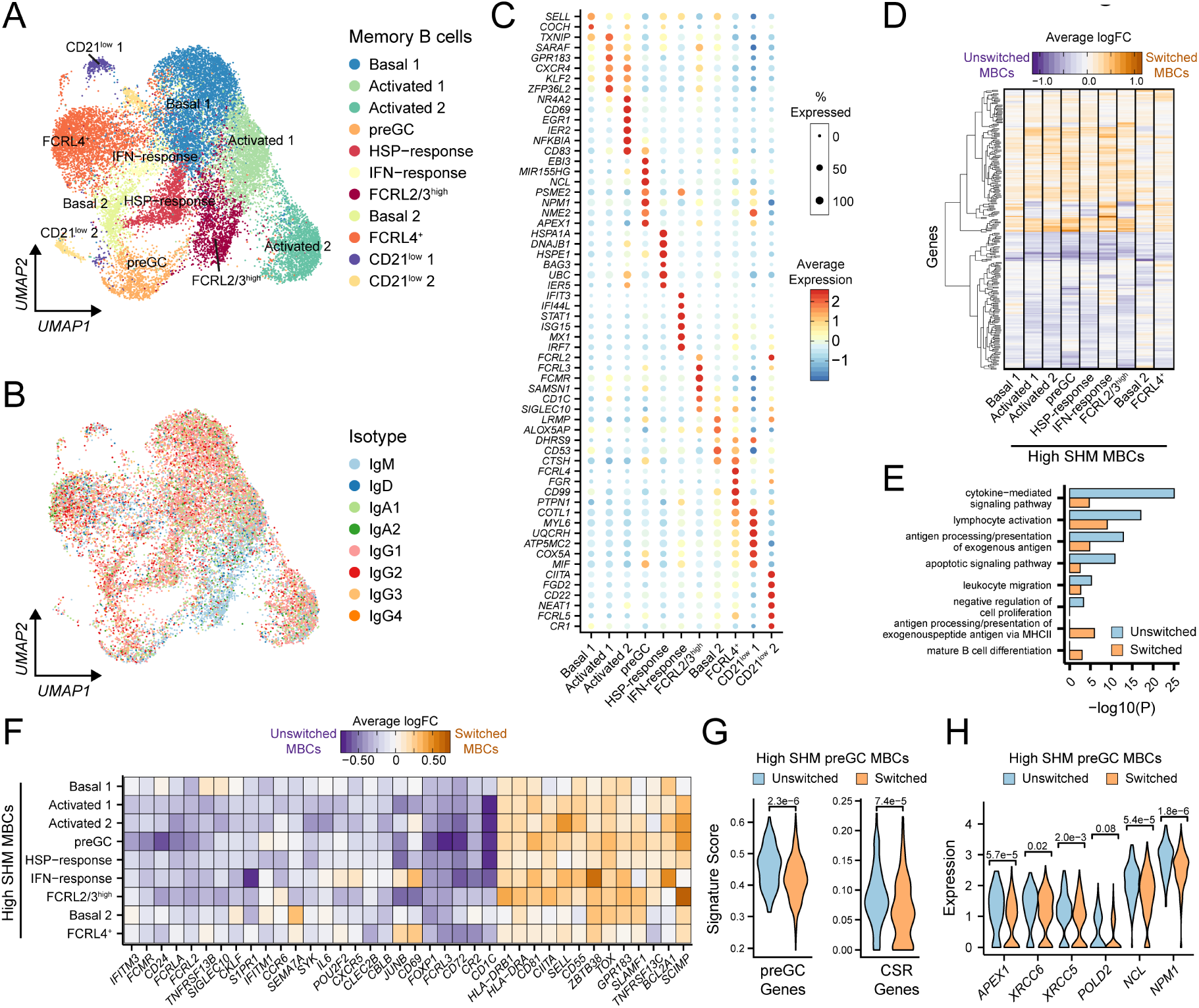
Diverse memory B cell states and antibody class-dependent gene expression. A) Clustering and UMAP visualisation of 21,595 memory B cell (MBC) single-cell transcriptomes. Identified cell populations include multiple basal and activated states, MBCs enriched for heat shock protein (HSP)-response and interferon (IFN)-response genes, preGC MBCs, FCRL2/3^high^ MBCs, tissue-resident FCRL4^+^ MBCs and two CD21^low^ populations. B) scVDJ-derived antibody isotypes of single MBC transcriptomes with high quality VDJ sequences (*n*=15,531 cells). C) Mean expression of top marker genes for MBC states. D) Average log fold change (logFC) of genes significantly enriched in switched or unswitched MBC states with similarly high affinity (based on SHM frequency). CD21^low^ clusters had too few cells and were excluded. E) Gene ontologies for genes significantly enriched in switched or unswitched MBCs with high affinity/SHM, in any cell state. F) Average logFC of selected immunologically-relevant genes significantly enriched in switched or unswitched MBC populations with high affinity/SHM. G) Single-cell scores for preGC and class switch recombination (CSR) signature gene sets in switched and unswitched preGC MBC with high affinity/SHM. H) Single-cell gene expression of key class switched recombination genes in switched and unswitched preGC MBC with high affinity/SHM.

We next considered whether class-switched and unswitched MBCs exhibit different gene expression networks that might reflect unique functional abilities (Dogan et al., 2009). We found little evidence that antibody class contributed towards the likelihood of an MBC to exist in a given state, with the exception of FCRL2/3^high^ MBCs which, similar to FCRL2/3^high^ GC B cells, were enriched for IgM^+^ cells (Figures 5B and S4F). Intriguingly, key marker genes associated with this state were broadly up-regulated in IgM^+^ cells across all MBC clusters (Figure S4G), suggesting a close relationship between the expression of these genes and IgM expression. We therefore compared gene expression of switched and unswitched MBCs of equivalent somatic hypermutation levels as a proxy for affinity (similar to our analysis of GC B cells in Figure 4), as MBCs expressing switched isotypes tended to have higher somatic hypermutation frequencies than unswitched MBCs (Figure S4H). We found widespread differences between unswitched and switched MBCs that were independent of MBC subset or state (Figure 5D), indicating a specific transcriptional wiring of switched and unswitched MBCs that might reflect altered abilities to signal to other immune cell populations, proliferate, survive, differentiate, migrate and respond to challenges (Figure 5E). In particular, we noted elevated expression in unswitched MBCs of genes known to regulate B cell migration within lymphoid tissues and genes with the potential to signal to and activate other immune cell types, including *IL6, CLEC2B, CR2* (CD21), *CKLF, S1PR1, CCR6*, and *SEMA7A* (Figure 5F; Arkatkar et al., 2017, Suzuki et al., 2008, Elgueta et al., 2015, Cinamon et al., 2004, Han et al., 2001, Welte et al., 2006). This could all contribute to the increased capacity of IgM^+^ MBCs to re-initiate GC reactions as part of a recall memory response (Dogan et al., 2009, Lutz et al., 2015, Seifert et al., 2015).

However, IgM^+^ MBCs were also enriched for many genes proposed to regulate or inhibit B cell activation, such as *FCRLA, FCRL2, FCRL3, CBLB, CD72* and *SIGLEC10* (Wu and Bondada, 2009, Sohn et al., 2003, Meyer et al., 2018, Kochi et al., 2009, Shabani et al., 2014), which may reflect a fine regulatory balance controlling their activation threshold. Perhaps underpinning this, unswitched MBCs also expressed higher levels of the transcription factor genes *POU2F2* (OCT2) and *FOXP1* than class-switched MBCs (Figure 5F), which coordinate the capacity of B cells to respond normally to antigen receptor signals and directly repress key regulators of plasma cell differentiation respectively (van Keimpema et al., 2015, Corcoran et al., 2014). This is consistent with switched IgG^+^ MBCs being more likely to differentiate into plasma cells, while unswitched IgM^+^ MBCs are more likely to re-enter or form secondary GC responses to gain higher affinity (Dogan et al., 2009, Lutz et al., 2015, Seifert et al., 2015). Indeed, we found that unswitched preGC MBCs exhibited significantly higher expression of many genes linked with the preGC state, including those associated with class switch recombination of their antibody genes (Figure 5G-H). This indicates that unswitched MBCs are more primed to undergo class switching for the first time as they re-enter the GC reaction, which will have an important impact on their subsequent selection dynamics in the GC.

## Discussion

Antibody responses are the foundation of effective immune memory and our ability to manipulate them through vaccination has contributed significantly to the success of modern medicine. While it is known that high affinity class-switched antibodies are generated during GC reactions, the complexity and dynamic nature of this response has presented a significant challenge for those seeking to understand how human B cell-mediated immunity is derived. By combining bulk antibody repertoire analysis with single-cell transcriptomics we have generated a detailed resource of the human GC response in a model secondary lymphoid tissue. This allowed us to define gene expression signatures of known and novel B cell states, most notably a population primed to undergo class switch recombination before entering the GC reaction. Whether a B cell undergoes class switch recombination at this stage then influences their capacity to undergo antibody-based selection within the GC and secondary activation as MBCs.

Although first histologically observed over a century ago, many questions remain about how B cells enter, experience and exit the GC reaction (Mesin et al., 2016, Shlomchik et al., 2019, Cyster and Allen, 2019). The practical challenges of sequentially sampling patient lymphoid tissues during an immune response make this especially true for understanding the most dynamic aspects of human B cell maturation. In particular, understanding the early events that facilitate GC entry by human B cells could provide new targets for adjuvants during vaccination or other immunotherapies. We have used pseudotemporal ordering to map the gene expression dynamics of both the early stages of B cell activation that correspond to antigen-dependent signalling through the BCR and the subsequent transition to a transcriptionally distinct preGC state, the latter of which is presumably under the regulation of cognate antigen-specific T helper cells. Our discovery that this preGC state is primed to undergo class switch recombination supports mounting evidence that class switching occurs before the classical GC response (Toellner et al., 1996, Roco et al., 2019, Pape et al., 2003) and has profound implications for understanding antibody-based selection dynamics in the GC.

Within the GC, B cell survival and selection is dependent on antigen binding to the BCR and its downstream signalling pathways (Kwak et al., 2019). By using single-cell transcriptomics paired with BCR sequence analysis we have been able to uncouple antibody class, affinity and B cell phenotype at single-cell resolution. We show that whether a B cell has undergone class switch recombination during GC entry is a major determinant of that B cell’s capacity to expand, acquire high antigen affinity and differentiate into plasmablasts or MBCs. IgG^+^ and IgA^+^ GC B cells have gene expression patterns consistent with increased BCR signalling and a greater capacity to remain within the GC, acquire T cell help and undergo somatic hypermutation to increase their affinity than GC B cells that have retained IgM expression. If class switch recombination does indeed occur predominantly prior to GC entry, as we and others suggest, these data support a model whereby the ability of a B cell to acquire high affinity is primarily dictated by the outcome of a specific class switch recombination “checkpoint” at the preGC stage. This would explain our observation that switched MBCs have comparable somatic hypermutation frequencies to switched plasmablasts, in contrast to the prevailing paradigm that higher affinity GC B cells preferentially differentiate towards the plasmablast fate whereas lower affinity clones seed the memory compartment (Suan et al., 2017b, Phan et al., 2006, Shinnakasu et al., 2016). The differences in affinity between these populations may instead be explained by the likelihood of whether they retained IgM expression prior to entering affinity maturation in the GC. This class switch recombination “checkpoint” is also relevant during the secondary activation of MBCs, where IgM^+^ MBCs appear more likely to undergo class switching compared to IgG^+^ or IgA^+^ MBCs, which may be important to provide a higher affinity secondary response through more prolonged GC maturation as well as more diverse effector functions from class-switched antibodies. Finally, our discovery of a conserved “unswitched” signature of elevated FCRL2 and FCRL3 expression (amongst other genes) across different B cell states, raises interesting questions about how these genes might regulate B cell function and immune responses more broadly.

Although the direct mechanisms shaping these gene expression differences between class-specific B cells remain to be elucidated, variations in the immunoglobulin tail tyrosine domain, linker flexibility or glycosylation sites between IgM and other antibody isotypes may all contribute to differential BCR signalling that could shape the behaviour of class-specific B cells (Martin and Goodnow, 2002, Engels et al., 2014, Xu et al., 2014b). Of note, we did not identify many major or meaningful differences in gene expression between IgG^+^ and IgA^+^ B cells, or between subclass-specific B cells, which may reflect the need to increase the power of future studies to identify potentially subtle gene expression differences between the less abundant isotypes. Unfortunately, we did not identify any IgE^+^ GC B cells, which are very rare in human tonsils, likely as they have been described to rapidly exit GCs due to IgE-specific BCR signalling (Haniuda et al., 2016). Given their relative availability, human tonsils are a useful tissue with which to examine the GC response. It will be of interest for future studies to compare class-specific gene expression differences in other tissues and contexts to determine the contribution of the local cellular environment on B cell maturation (James et al., 2020, Smillie et al., 2019). Similarly, recent technical advances allowing the simultaneous readout of antigen specificity, antibody sequences and gene expression (Setliff et al., 2019) make it possible to examine how different types of antigens may be linked with different B cell phenotypes. Finally, our profiling of human MBCs in a secondary lymphoid tissue revealed diverse states reflecting different activation, signalling and functional potential. Given an emerging appreciation for heterogeneity within both human and mouse MBC populations (Good-Jacobson and Shlomchik, 2010), our single-cell characterisation of different MBC states will act as a valuable resource to interrogate the potential relevance for such diverse populations in mediating humoral immunity.

Together, our integrated analyses of antibody repertories and gene expression of human B cell states during affinity maturation highlight how the outcome of class switch recombination is a major determinant of B cell fate and function. More broadly, our detailed annotation of diverse B cell states provides a new and uniquely detailed framework through which to view B cell-mediated immune responses in the context of both health and B cell-related pathologies such as allergy, multiple sclerosis, rheumatoid arthritis and lymphoma.

## Methods

### Human ethics, tissue collection and sorting of B cell subsets

Routine tonsillectomy patients at the Royal London Hospital aged between 3 and 14 were consented for tissue collection with approval from North West - Greater Manchester East Research Ethics Committee under REC reference 17/NW/0664. Following removal of dead tissue and clotted blood, each palatine tonsil was bisected and processed separately as follows, with all repertoire and single-cell analyses performed on a single bisected sample. Tonsillar tissue was dissected manually into approximately 2-3 mm pieces prior to homogenisation with the gentleMACS™ Dissociator using C tubes and two rounds of the Multi_C_01_01 setting in 8 mL RPMI + 10% fetal calf serum (FCS). Dissociated cells were then passed through a 70 μM filter prior to isolation of mononuclear lymphocytes using Ficoll-Paque™ gradients. Isolated mononuclear cells were then washed in RPMI + 10% FCS before cell counting and viability determination using Trypan Blue staining. Cells to be used for 10X single-cell transcriptomics were processed immediately, while cells for bulk repertoire sequencing were either stored overnight at 4°C or cryopreserved in FCS with 10% DMSO at −70°C.

For bulk B cell repertoires, we labelled 1-1.3×10^8^ tonsillar lymphocytes per donor for fluorescence-activated cell sorting (FACS). Briefly, cells were washed and stained with Zombie NIR™ Fixable Viability Kit (BioLegend) to label dead cells, followed by washing with FACS buffer (PBS + 0.5 % BSA + 2 mM EDTA) and incubation with human FcR Blocking Reagent (Miltentyi Biotec). Cells were then stained with CD19-APC (clone HIB19; BioLegend), CD38-PE-Cy7 (clone HB-7; BioLegend), CD27-Pacific Blue™ (clone O323; BioLegend), IgD-PerCP-Cy5.5 (clone IA6-2; BioLegend), and IgM-FITC (clone MHM-88; BioLegend). For bulk B cell repertoires, two aliquots of 250,000 cells for the following populations were sorted using a BD FACSAria™ IIIu: total B cells (live CD19^+^), naïve B cells (live CD19^+^ IgD^+^ CD38^-^), germinal centre B cells (live CD19^+^ IgD^-^ CD38^+^), memory B cells (live CD19^+^ IgD^-^ CD38^-^), and plasmablasts (live CD19^+^ IgD^-^ CD38^++^). Gates were set using fluorescence minus one (FMO) controls. Sorted B cell samples were processed immediately for RNA extraction. For single-cell RNA-seq, 50,000-200,000 class-switched memory B cells (live CD19^+^ IgD^-^ CD38^-^ IgM^+^ (*n*=2) or IgM^-^ (*n*=4)) were sorted.

### Bulk VDJ repertoire library preparation and sequencing

RNA was isolated from sorted B cell aliquots using the RNAqueous™-Micro Total RNA Isolation Kit (ThermoScientific) supplemented with β-mercaptoethanol according to manufacturer’s protocol. RNA was stored long-term at −80°C or processed immediately to generate bulk repertoire sequencing libraries of immunoglobulin heavy chains (IgH) as previously described (Horns et al., 2016), with minor changes. Briefly, 50 to 100 ng RNA from sorted B cell subsets were annealed to a pooled set of five isotype-specific IgH constant region primers containing unique molecular identifiers (UMIs) of either 10 or 12 nucleotides at 72°C for 5 minutes before being immediately placed on ice for 2 minutes. We then performed first-strand cDNA synthesis using SuperScript IV reverse transcriptase (ThermoFisher Scientific) with recommended reagent concentrations and the following cycling conditions in a thermocycler: 105°C lid; 55°C 10 minutes; 80°C 10 minutes; 4°C hold. Second-strand cDNA synthesis was performed using Phusion® High-Fidelity DNA Polymerase (NEB) and six IgH variable region primers containing 10 or 12 nucleotide UMIs with the following cycling conditions: 105°C lid; 98°C 4 minutes; 52°C 1 minutes; 72°C 5 minutes; 4°C hold. Double-stranded cDNA was then purified using (0.6X) Ampure XP beads (Beckman Coulter) before amplification with Illumina adapter-containing primers (Nextera i7 indices) and NEBNext Ultra II Q5 Master Mix (NEB) as follows: 105°C lid; 98°C 30 seconds; (98°C 10 seconds, 72°C 50 seconds) × 22 to 28 cycles; 72°C 2 minutes; 4°C hold. Amplified libraries were purified using (0.6X) Ampure XP beads and quantified by Qubit™ dsDNA HS Assay Kit prior to multiplexing. Libraries were sequenced with PhiX spike-in using paired-end 301 bp reads on the Illumina MiSeq platform.

### 10x Genomics Chromium single-cell transcriptomics and VDJ library preparation, sequencing and raw data processing

Total tonsillar immune cells (*n*=7) or FACS-enriched memory B cells (*n*=6) were loaded according to the manufacturer’s protocol for either for the Chromium single-cell 3’ kit (v2; n = 1) or 5’ gene expression (v1; n = 6) to attain between 2000-5000 cells per well. Library preparation for both gene expression and VDJ (BCR) was performed according to the manufacturer’s protocol prior to sequencing on the Illumina NextSeq 500 platform with 26/8/134 bp or 155/8/155 bp read configurations respectively. BaseCall files were subsequently used to generate library-specific FASTQ files with cellranger mkfastq (v3.0.0) prior to running cellranger count (v3.0.0) with the GRCh38 (release 92) reference to produce cell barcode-gene expression matrices using default settings. For single-cell VDJ datasets, cellranger vdj (v3.0.0) was run using the refdata-cellranger-vdj-GRCh38-alts-ensembl-2.0.0 reference from 10x Genomics using default settings. Poor quality contigs that either did not map to immunoglobulin chains or were designated incomplete by cellranger were discarded. We further filtered IgH contigs as to whether they had sufficient coverage of constant regions to ensure accurate isotype assignment between closely related subclasses using MaskPrimers.py (pRESTO v0.5.10; Vander Heiden et al., 2014).

### Quality control and sequence assembly of bulk B cell repertoires

Raw sequencing data from bulk VDJ libraries were processed to generate UMI-collapsed consensus VDJ sequences using pRESTO (v0.5.10; Vander Heiden et al., 2014). Paired-end sequencing reads with mean Phred quality scores less than 25 were removed, and remaining sequences were annotated and trimmed for PCR primer and UMI sequences. UMI barcodes were then filtered by length and the presence of ambiguous nucleotides, prior to UMI alignment using MUSCLE (v3.8.31; Edgar, 2004). UMI consensus sequences were then generated, with a minimum of three reads per UMI required, prior to assembly of paired-end UMI consensus sequences into a single VDJ contig and annotation of constant region isotype using MaskPrimers.py align (v0.5.10; Vander Heiden et al., 2014). Duplicate VDJ sequences within each subset were then collapsed using CollapseSeq.py (v0.5.10; Vander Heiden et al., 2014) before VDJ gene assignment and functional annotation using AssignGenes.py (ChangeO v0.4.5; Gupta et al., 2015) and IgBLAST (v1.12.0; Ye et al., 2013).

### Identification of clonally-related sequences, genotype inference and calculation of IgH mutation frequencies

Following initial quality control, all single-cell VDJ sequences were combined together with bulk BCR repertoire sequences from the same donor for subsequent processing. IgH sequences were annotated using IgBLAST (v1.12.0; Ye et al., 2013) and assigned isotype classes using AssignGenes.py prior to correction of ambiguous V gene assignments using TIgGER (v0.3.1; Gupta et al., 2015, Gadala-Maria et al., 2015). Clonally-related IgH sequences were identified using DefineClones.py (ChangeO v0.4.5; Gupta et al., 2015) with a nearest neighbour distance threshold of 0.0818, as determined by the mean 99% confidence interval of all 8 donors with distToNearest (Shazam v0.1.11; Gupta et al., 2015). CreateGermlines.py (ChangeO v0.4.5) was then used to infer germline sequences for each clonal family and observedMutations (Shazam v0.1.11) was used to calculate somatic hypermutation frequencies for each IgH sequence. Sequences with mutation frequencies greater than 0.02 were annotated as “High” mutation levels, those between 0 and 0.02 as “Low” mutation levels and 0 as “None”. For bulk BCR repertoire analysis in Figure 1, single-cell VDJ sequences were excluded and analysed, providing ∼1.5 million high-confidence and unique IgH sequences, with a median of 14 UMIs per sequence, a median of 28,918 unique sequences per donor per subset and approximately 96-99% of these sequences appeared to be functional.

Single-cell VDJ analysis was performed broadly as described previously (James et al., 2020). Briefly, the number of quality filtered and annotated IgH, IgK or IgL were determined per unique cell barcode prior to integration with single-cell gene expression objects. If more than one contig per chain was identified, metadata for that cell was ascribed as “Multi”. IgH diversity analyses were performed using the rarefyDiversity and testDiversity of Alakazam (v0.2.11; Gupta et al., 2015). To assess clonal relationships between cell types, co-occurrence of expanded clone members between cell types was reported as a binary event for each clone that contained a member within two different cell types in either single-cell or bulk repertoires. For comparisons of somatic hypermutation and isotype frequencies between subsets we used Wilcoxon Rank Sum Signed test while Student’s t test was used to compare mean clonal diversity scores.

### Data quality control, processing and annotation of single-cell RNA-seq

Gene expression count matrices from cellranger were used to calculate percentage mitochondrial expression per cell barcode prior to mitochondrial genes being removed from gene expression matrices. Similarly, the V, D and J gene counts for each immunoglobulin and T cell receptor were summed to calculate an overall expression before individual genes were removed from gene expression matrices. Counts of individual IgH constant region genes were also summed together (IgG1-4, and IgA1-A2) and removed from gene expression matrices. Modified gene-by-cell matrices were then used to create Seurat objects for each sample using Seurat (v3.0.3; Butler et al., 2018, Stuart et al., 2019), removing genes expressed in fewer than 3 cells. Cell barcodes with >1000 and <60000 UMIs and >500 and <7000 genes detected were removed, as were cell barcodes with >30% mitochondrial reads. Individual samples were then log transformed, normalised by a factor of 10000 prior to predicting cell cycle phases using the CellCycleScoring command and then identifying the 3000 most variable genes within each sample using the “vst” method. We then performed a preliminary integration of all unsorted immune cells or all sorted memory B cell datasets together using FindIntegrationAnchors and IntegrateData (3000 genes) before regressing out cell cycle scores and mitochondrial gene expression, performing principle component analysis (PCA) and preliminary clustering and cell type annotation. One cluster was identified to be enriched with predicted doublets based on the results from DoubletFinder (v2.0.1; McGinnis et al., 2019) and scrublet (v0.2; Wolock et al., 2019), and a small number of cell barcodes with co-expression of B/T/non-lymphoid markers were manually removed by filtering on UMAP coordinates. Following the removal of poor quality cell barcodes from gene expression matrices based on these preliminary analyses of the unsorted immune cell and sorted memory B cell libraries, we then integrated all normalised count matrices together using the unsorted immune cell count matrices as a reference with 4000 highly variable genes before scaling the integrated data and regressing cell cycle and mitochondrial gene expression, running PCA and identifying broad cell type lineages (B cell, T cell and non-lymphoid cells) using a broad resolution for clustering. These lineages were then separated for more detailed cell state annotation by recomputing the PCA (RunPCA), nearest neighbour graph (FindNeighbors) and unbiased clustering (FindClusters). Uniform Manifold Approximation and Projection (UMAP) was then used to visualise both integrated and lineage-specific datasets. B cells were annotated with scVDJ metadata from the integrated repertoire analysis detailed above.

### Differential gene expression and signature enrichment analysis

Gene expression markers for different clusters of unsorted B cells, T cells, non-lymphoid cells and sorted MBCs were identified using FindAllMarkers from Seurat with default settings, including Wilcoxon test and Bonferroni *p* value correction (v3.0.3; Butler et al., 2018, Stuart et al., 2019). Differential gene expression for antibody class-specific or somatic hypermutation frequency for GC B cells or class-switched MBCs was performed using FindAllMarkers with Benjamini-Hochberg false discovery rate (FDR) correction. Genes were deemed significantly different if FDR < 0.05, average log fold change > 0.1 and the gene was detected in >20% of cells in that group. Gene ontology analyses for high affinity/SHM class-specific GC B cells were performed with Ingenuity Pathway Analysis (Qiagen) software using avg_logFC values of all genes significantly enriched in at least one class. For IFN-response MBC cluster gene ontology enrichment was performed with all significant gene markers for this cluster in Metascape (Zhou et al., 2019) using default settings, as were all genes significantly enriched or depleted in switched or unswitched in at least one MBC cluster.

Enrichment of gene set signatures for single cells was calculated using AUCell (v1.5.5; Aibar et al., 2017). For class switch recombination, a manually curated shortlist of genes was determined for genes linked with CSR that were reliably detected in our sparse scRNA-seq datasets (Stavnezer and Schrader, 2014). Gene signatures of high and low affinity mouse LZ GC B cells (Shinnakasu et al., 2016) were obtained by quantifying RNA-seq transcript counts against GRCm38 transcriptome build using Salmon (v1.0.0; Patro et al., 2017), collapsing protein-coding transcripts into a single gene count using tximport (v1.10.1; Soneson et al., 2016), identifying significant gene expression differences between the two groups using DESeq2 (v1.22.2; Love et al., 2014) with a threshold of log fold change > 1.5 and *padj* < 0.05 and converting mouse gene IDs to human using bioMart (v2.38.0; Durinck et al., 2005, Durinck et al., 2009). For IgM stimulation gene sets, Geo2R (Barrett et al., 2012) was used to analyse microarray data from a timecourse of wild type splenic mouse B cells stimulated with 10 µg/ml of anti-IgM (Shinohara et al., 2014) and identify genes enriched following α-IgM treatment compared to control untreated cells with FDR < 0.05 and a fold change > 2. To calculate preGC and FCRL2/3^high^ signature scores in MBC subsets, the top 50 most significantly enriched gene markers per cluster in the unsorted B cell subset analysis were used with AUCell. Unless indicated otherwise, Wilcoxon Ranked Signed Sum test was used to test for significant differences.

### Prediction of cell-cell communication using CellPhoneDB

To evaluate potential cell-cell communication, we used CellPhoneDB (v2.0.6; Vento-Tormo et al., 2018, Efremova et al., 2020) to examine the expression of ligand-receptor pairs between different scRNA-seq clusters. Briefly, we exported raw gene count matrices from Seurat, converted gene IDs to Ensembl IDs using bioMart (v2.38.0; Durinck et al., 2005). We re-annotated all non-lymphoid cell type clusters as antigen-presenting cells (APCs), naïve and effector T cell groups by CD4 or CD8 expression, Treg and Tfr as “Treg” and rare GC subsets (prePB and FCRL2/3^high^) as “GC” and exported cell type metadata for use with raw count data using the “statistical_analysis” command of CellPhoneDB with database v2.0.0. The number of unique significant ligand-receptor co-expression pairs (putative interactions; *p* value < 0.05) between each cell type was then counted and visualised as a heatmap, while exemplar interacting pairs were visualised by calculating mean average expression level of gene 1 in cell type 1 and gene 2 in cell type 2 are indicated by colour and *p* values indicated by circle size.

### RNA velocity and pseudotemporal ordering

To calculate single-cell velocities we first quantified spliced and unspliced transcripts for the filtered barcodes output from cellranger using velocyto (v0.17.10; La Manno et al., 2018). Loom files were then combined using loompy (v2.0.17) before reformatting cell barcode names to be compatible with Seurat objects and merging with a Scanpy (v1.4; Wolf et al., 2018) object containing the raw gene expression matrix of high quality annotated single B cell transcriptomes (see above) using scVelo (v0.1.23; Bergen et al., 2019). scVelo was then used to pre-process, filter and normalise velocyto-derived counts with default settings prior to computation of the first- and second-order moments (scv.pp.moments) and subsequent velocity estimation using a dynamical model (scv.tl.recover_dynamics and scv.tl.velocity). Velocities were then projected and visualised onto UMAP embeddings at a grid level using an inverted transition matrix obtained from scv.tl.transition_matrix prior to scv.tl.velocity_embedding, basis=‘umap’.

For pseudotemporal ordering of the B cell activation and GC entry trajectory, a partition-based graph abstraction (PAGA) was performed for Naïve, Activated, preGC and LZ GC B cell clusters (Scanpy; tl.paga) before computing connectivity of single cells using a diffusion map (Scanpy; tl.diffmap). Velocity-based pseudotime reconstruction was performed using default settings for scVelo commands tl.recover_latent_time and tl.velocity_pseudotime, although the pseudotemporal order was reversed to place naïve cluster at pseudotime = 0. Dynamic gene expression changes were examined by using tl.rank_velocity_genes (scVelo) to sub-cluster original cell type annotations (resolution = 1) based on RNA velocity and the top 200 genes per sub-cluster were reported before filtering out ribosomal genes and collapsing to unique genes. Genes were then clustered through pseudotime for heatmap visualisation with smoothed expression scores in scVelo. Quantitation of individual gene expression or AUCell-derived signature scores of single-cells across pseudotime was performed using smoothed normalised counts with geom_smooth() including 95% confidence intervals. For pseudotemporal analysis of GC B cell subsets, a similar approach was taken for the LZ GC, GC, DZ GC, FCRL2/3^high^ GC and prePB clusters, except that diffusion-based pseudotime was calculated with Scanpy (tl.dpt) independent of RNA velocity measurements. Visualisation of a custom list of top GC subset marker genes for LZ GC, GC and DZ GC clusters was performed using pl.paga.path heatmap with Scanpy.

### Immunohistochemistry

Tonsil biopsies were cut from formalin fixed paraffin-embedded blocks then deparaffinized in xylene and rehydrated through a series of ethanols to water. Endogenous peroxidase was blocked with 3% hydrogen peroxide. Heat-mediated antigen retrieval was performed using a commercial citrate-based unmasking buffer (Vector Labs) at 120°C using a pressure cooker. Sections (3µm thick) were then incubated for 40 minutes at RT with 1:1000 dilution of anti-APE1 (HPA002564; Sigma). Detection of primary antibody was performed using the Super-sensitive–Polymer HRP system (Biogenex) and staining visualized using purple chromogen VIP (Vector Labs) and hematoxylin as a nuclear counterstain. Slides were then scanned (Pannoramic 250 Flash) before being left to soak in xylene to de-coverslip. Once the coverslips were removed, slides were rehydrated through ethanol to water. De-staining and stripping of the primary antibodies and the heat-labile VIP chromogen was achieved using a subsequent round of heat-mediated antigen retrieval as per the first round of staining. The second primary antibody CD20 (M0755; Dako) was incubated for 40 minutes at RT at a dilution of 1:500, followed by detection, visualization and scanning as before. Negative controls were performed by treating sequential sections as above but without the second primary antibody (CD20) to confirm complete antibody and signal stripping. Images were prepared using CaseViewer (3DHistTech).

### Quantitation of IgH germline transcripts

To quantify expression of IgH germline transcripts (GLT), all mapped reads to the IgH locus (chr14:105540180-105879151) were extracted from cellranger-derived bam files. DropEst (v0.8.6; Petukhov et al., 2018) was then used to count reads against a custom GTF containing coordinates for I promoter GLT sequences (Sideras et al., 1989, Fujieda et al., 1996), annotation of membrane-specific IgH exons and IgH switch regions identified by enrichment of the WRCY motif using HOMER2 (v4.9.1; Heinz et al., 2010). Counts were then read into Seurat without filtering for log10-normalisation and scaling. Due to mapping ambiguities between subclass-specific regions, subclass-specific counts were summed together.

### Accession codes and data availability

Raw sequencing and processed data files for single-cell RNA sequencing, single-cell VDJ sequencing, and bulk B cell repertoires are available at ArrayExpress (accession numbers: E-MTAB-8999, E-MTAB-9003 and E-MTAB-9005). RNA-seq and microarray data for high affinity and α-IgM stimulation gene expression signatures were obtained from GSE73729 and GSE41176 respectively. DNase-seq and ChIP-seq datasets from ENCODE were visualised using UCSC Genome Browser (Kent et al., 2002, Rosenbloom et al., 2012).

## Acknowledgements

This research was supported by funding from the Wellcome Trust to L. K. J. (208961/Z/17/Z), S. A. T. (206194) and J. C. R. (110020/Z/15/Z). H. W. K. was funded by a Sir Henry Wellcome PostDoctoral Fellowship (213555/Z/18/Z). We would like to thank the Bart’s and the London Genome Centre at QMUL for library sequencing support. We also thank members of the James and Teichmann labs for their help and support, especially Mirjana Efremova for her help with CellPhoneDB. Finally, we are grateful to Neil McCarthy, Kylie James, Kerstin Meyer, Lou Herman and Jo Spencer for their help reviewing the manuscript.

## Author contributions

H. W. K. initiated the project, designed and performed experiments, analysed data and wrote the manuscript. N. O. carried out tonsillectomy tissue collection. J. C. R. and A. J. C. designed and performed immunohistochemistry experiments. G. W. assisted in FACS sorting. S. A. T. supervised and interpreted data analysis. L. K. J. initiated the project, designed and supervised experiments and data analysis, interpreted data and wrote the manuscript.

## Supplementary Figures

**Figure S1.**
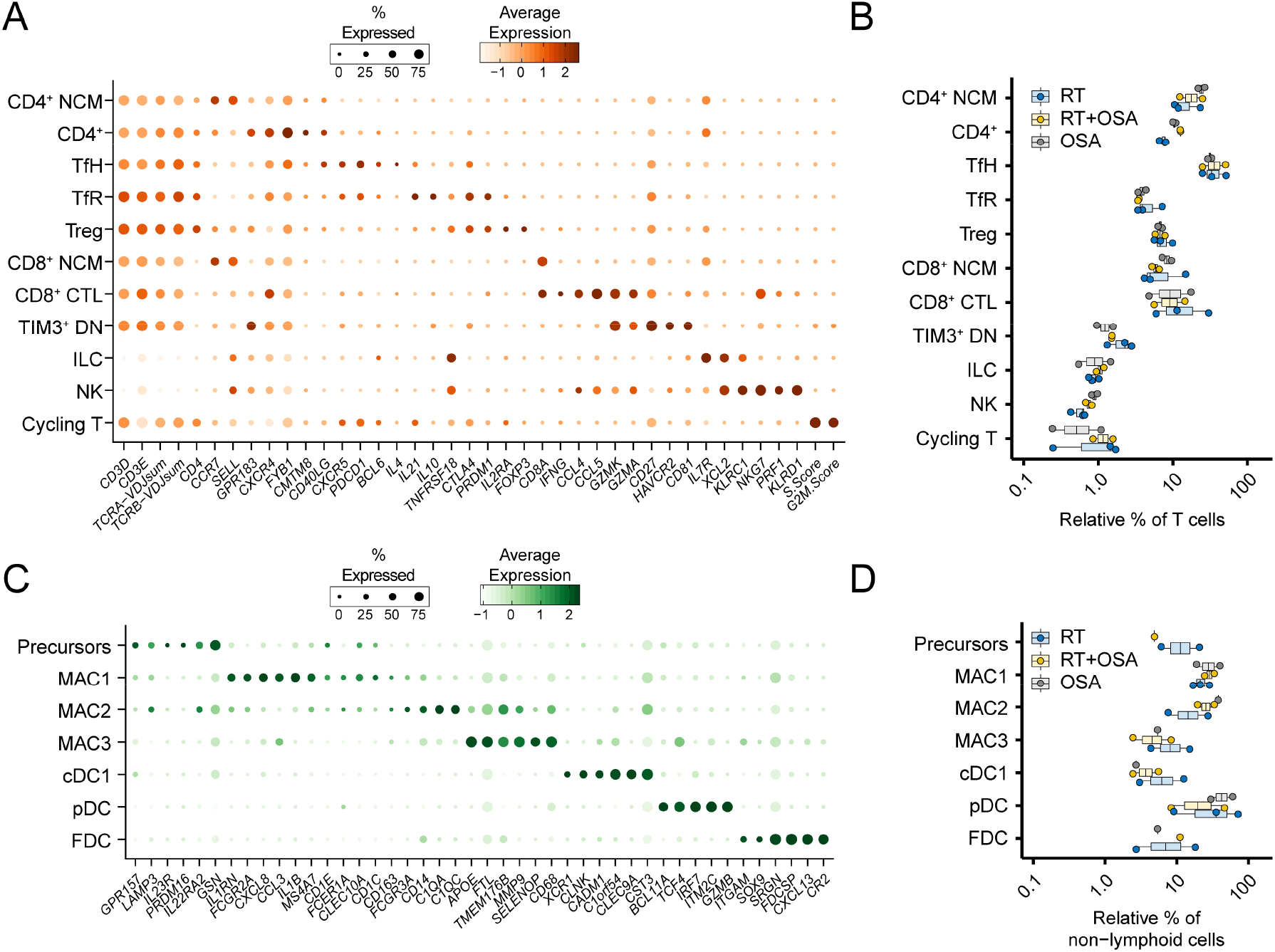
Annotation of non-B cell populations in the human tonsils. A) Mean expression of key marker genes used to define T cell scRNA-seq clusters, including CD4^+^ naïve or central memory (CD4^+^ NCM), CD4^+^, T follicular helper (TfH), T follicular regulatory (TfH), T regulatory (Treg), CD8^+^ naïve or central memory (CD8^+^ NCM), CD8^+^ cytotoxic (CD8^+^ CTL), TIM3^+^ CD4/CD8 double-negative (TIM3^+^ DN) and cycling T cells, in addition to innate lymphoid cells (ILC) and natural killer (NK) cells. Frequency of cells for which each gene is detected is denoted by size of the dots. B) Relative frequencies of different T cell subsets separated by clinical indication for tonsillectomy. OSA; obstructive sleep apnoea (*n* = 2), RT; recurrent tonsillitis (*n* = 3), RT+OSA (*n* = 2). C) Mean expression of key marker genes used to define non-lymphoid cell scRNA-seq clusters, including monocyte/macrophages precursor (Precursors), macrophage (MAC1, MAC2, MAC3), conventional dendritic cell 1 (cDC1), plasmacytoid-derived dendritic cell (pDC) and follicular dendritic cell (FDC) subsets. Frequency of cells for which each gene is detected is denoted by size of the dots. D) Relative frequencies of different non-lymphoid cell subsets separated by clinical indication for tonsillectomy.

**Figure S2.**
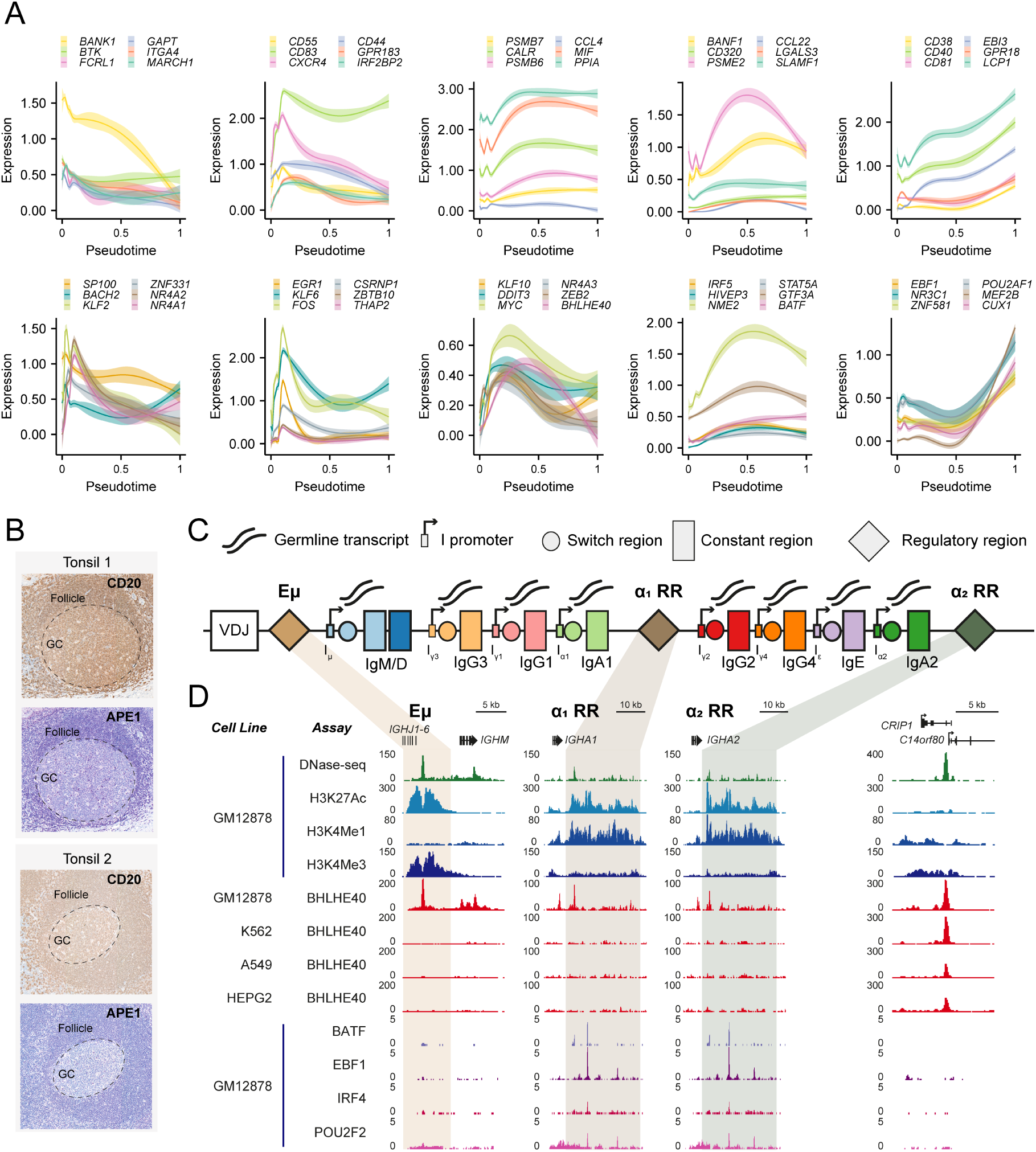
Dynamic gene expression during class switch recombination and transcription factor binding at the immunoglobulin locus. A) Smoothed gene expression for example cell surface receptor or cytokine (top row) and transcription factor (bottom row) genes that are differentially regulated through velocity-based pseudotime of B cell activation and GC entry. B) Immunohistochemistry of CD20 (B cell marker) and APE1 (*APEX1*) in two paediatric human tonsils reveals depleted expression of APE1 in germinal centres (GCs) compared to the follicular zone. C) Schematic of the human immunoglobulin heavy chain (IgH) locus, with intergenic (I) promoters, switch regions, germline transcripts and regulatory regions (Eμ, α_1_ RR, α_2_ RR). D) Open chromatin (DNase-seq) and ChIP-seq from ENCODE consortium at Eμ, α_1_ RR, α_2_ RR, and a control neighbouring locus (*CRIP1 / C14orf80*) for EBV-transformed B lymphocyte cell line GM12878 and control non-B lymphocyte cell lines (K562, A549, HEPG2).

**Figure S3.**
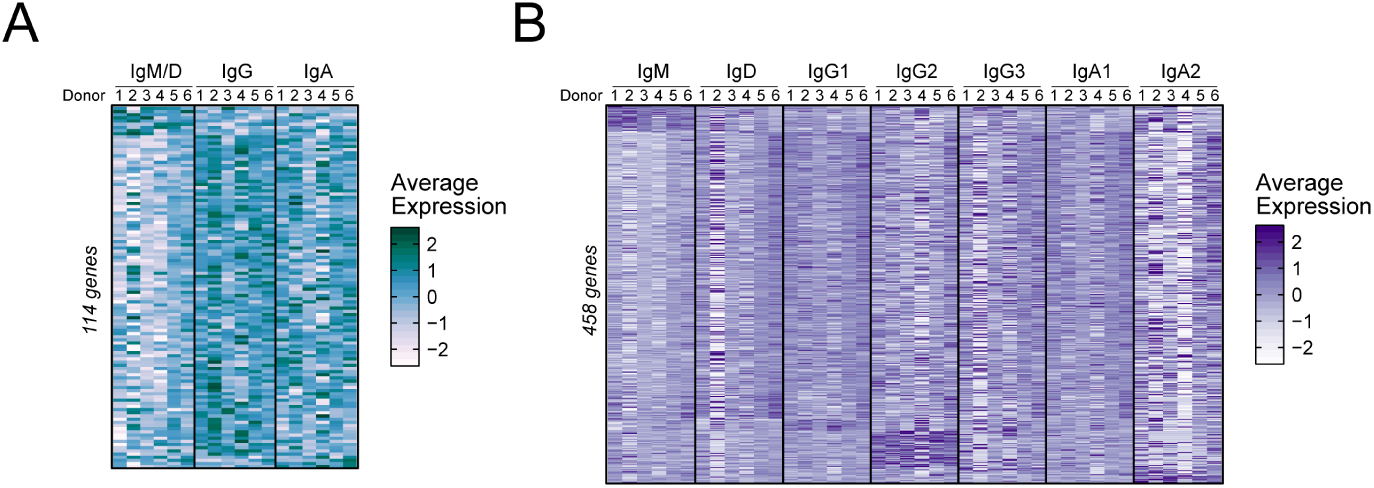
Class- and subclass-specific gene expression analyses. A) Pseudobulk heatmaps of average expression per donor of differentially expressed genes between class-specific GC B cells with similar affinity (based on SHM frequency). B) Same as in A), but for subclass-specific gene expression analyses.

**Figure S4.**
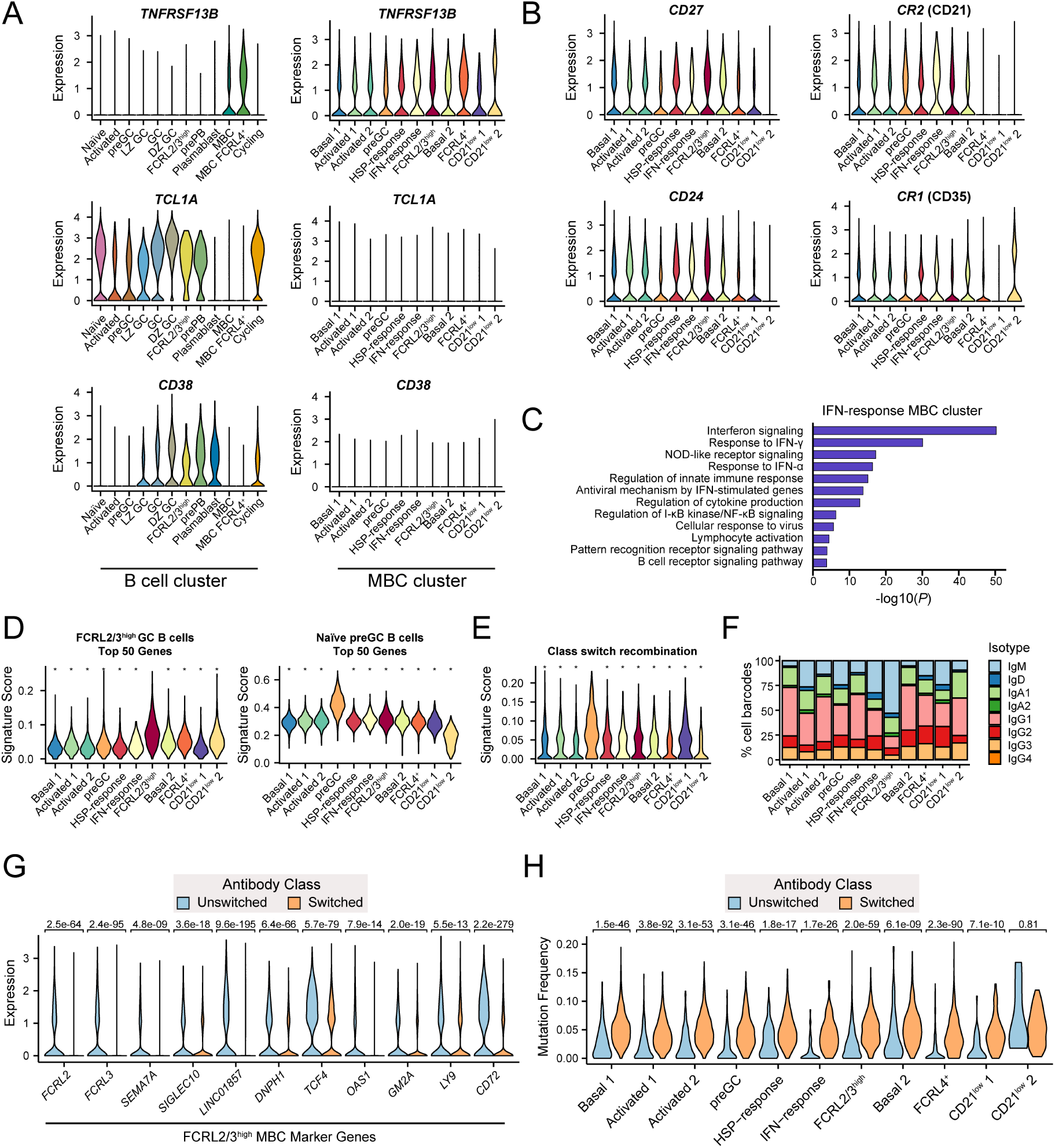
Characterisation of memory B cell states identified by scRNA-seq. A) Single-cell expression of memory B cell (*TNFRSF1B*), naïve/undifferentiated (*TCL1A*) and germinal centre (*CD38*) markers across all B cell subsets (left) and sorted memory B cell subsets (right). B) Single-cell expression of key marker genes differentially expressed by CD21^low^ MBC populations. C) Top gene ontologies for significantly enriched genes in the IFN-response MBC cluster. D) Single-cell AUCell-derived scores for top 50 marker genes of the naïve preGC B cells and FCRL2/3^high^ GC B cells in MBC subsets. * denotes *p* value < 0.001. E) Single-cell AUCell-derived scores for class switch recombination gene set in MBC subsets. * denotes *p* value < 0.001. F) Relative frequencies of scVDJ-derived antibody subclass expression within different MBC scRNA-seq populations. G) Single-cell expression of key marker genes of the FCRL2/3^high^ B cell states between switched and unswitched MBCs. H) Somatic hypermutation frequencies of scVDJ-derived antibody genes between switched and unswitched B cells within different MBC populations.

